# The GA4GH Workflow Execution Service: A Standard API for Workflow Execution and Management in Genomics

**DOI:** 10.64898/2026.10.20.760340

**Authors:** Valentin Schneider-Lunitz, Landfried Kraatz, Patrick Magee, Venkat S. Malladi, Brian D. O’Connor, Susheel Varma, James Eddy, Hirotaka Suetake, Jeff Gentry, Michael R. Crusoe, Angel Pizarro, Junjun Zhang, Denis Yuen, Kyle Ellrott, Philip R. Kensche, Michael Lukowski, Reggan Thomas, Sasha V. Siegel, Alexander Kanitz, Tazro Ohta, Sven O. Twardziok

**Affiliations:** Berlin Institute of Health at Charité – Universitätsmedizin Berlin, Center of Digital Health, Charitéplatz 1, 10117 Berlin, Germany; Grafana Labs, New York, NY 10006, USA; Verily Life Sciences, Redwood City, CA, USA; Nimbus Informatics, Kenmore, WA, USA; Sage Bionetworks, Seattle, WA, USA; University of Sheffield, Sheffield, UK; Fred Hutch Cancer Center, Seattle, WA, USA; Sator, Inc., Tokyo, Japan; Fulcrum Genomics, Somerville, Massachusetts, USA; Freie Universität Berlin, Germany; NVIDIA, Santa Clara, CA, USA; Adela Inc, Foster City, CA, USA; Adaptive Oncology, Ontario Institute for Cancer Research, Toronto, Ontario M5V 3S1, Canada; Department of BioMedical Engineering, Oregon Health & Science University, 3181 SW Sam Jackson Park Rd, Portland, OR, 97239, USA; German Cancer Research Center (DKFZ) Heidelberg, Division Omics IT and Data Management Core Facility (ODCF), Im Neuenheimer Feld 280, 69120 Heidelberg, Germany; University of Chicago, Chicago, IL, USA; Global Alliance for Genomics and Health (GA4GH),Toronto, ON, Canada; European Molecular Biology Laboratory, European Bioinformatics Institute (EMBL-EBI), Hinxton, UK; Biozentrum, University of Basel, Spitalstrasse 41, 4056 Basel, Switzerland; Swiss Institute of Bioinformatics, Amphipôle, Quartier UNIL-Sorge, 1015 Lausanne, Switzerland; Database Division for Life Science (DBCLS), BioData Science Initiative (BSI), National Institute of Genetics, Research Organization of Information and Systems, Japan; German Center for Child and Adolescent Health (DZKJ), partner site Berlin, Berlin, Germany; German Center for Mental Health (DZPG), partner site Berlin/Potsdam, Germany

## Abstract

**Motivation:** Rapid growth in life science data generation has led to large and increasingly distributed collections of sensitive genomic data. To analyse this data, researchers increasingly rely on computational workflows to enable scalable, portable and reproducible analysis. However, these workflows are executed by specialized workflow engines across heterogeneous compute environments, each with its own requirements. As a result, users often need engine- and compute-environment-specific knowledge to submit, monitor and manage workflow execution, increasing the barriers to interoperability. These challenges are compounded in large-scale projects, where workflow runs must be dispatched to local clusters, cloud platforms or data hosting infrastructures with different underlying technologies.

**Results:** We describe the Global Alliance for Genomics and Health (GA4GH) Workflow Execution Service (WES) API, an approved GA4GH standard first released in 2018, together with its data model and the endpoints that define a unified interface for workflow execution. By standardizing workflow invocation, the WES API allows a single client to submit, monitor and cancel workflow runs across diverse workflow engines and compute backends, such as scheduling clusters or cloud platforms. We surveyed thirteen adoptions of the standard, covering standalone services and platforms in four programming languages. In a two-site demonstration, a WES implementation from Germany and one from Japan executed the same federated allele-frequency analysis with site-specific adjustments and only aggregated results leaving either site, demonstrating that it is possible to run the same analysis across independently built and operated WES services in different countries. From the gaps identified, we propose extensions for community discussion.

**Availability and implementations:** The latest version of the WES API repository can be found at https://github.com/ga4gh/workflow-execution-service-schemas (DOI: https://doi.org/10.59756/m382x-ww ). The full Reference Documentation can be viewed at: https://ga4gh.github.io/workflow-execution-service-schemas/docs/

## Introduction

Rapid developments in genomics, healthcare, and computational biology have led to an exponential growth in volume and diversity of decentralized datasets. Computational workflows, which define reproducible sequences of data processing and analysis steps, have become a cornerstone of modern data-driven research. By wrapping analytical procedures, software dependencies, and parameters into a structured pipeline, workflows enable automation, portability, scalability and reproducibility of computational analysis across different computing environments (Ahmed et al., 2021; Boettiger, 2015; Gruning et al., 2018; Kim et al., 2018; Wratten et al., 2021). To support the findability and reusability of workflows, Dockstore (Yuen et al., 2021) and WorkflowHub (Gustafsson et al., 2025) provide community platforms where researchers can discover and publish workflows described using community-adopted workflow languages (Amstutz et al., 2016; Di Tommaso et al., 2017; Koster & Rahmann, 2012; Voss K, 2017), engines (Di Tommaso et al., 2017; Koster & Rahmann, 2012; Vivian et al., 2017; Voss K, 2017) and community-curated pipeline collections (Ewels et al., 2020). These resources play an important role in promoting the FAIR (**F**indable, **A**ccessible, **I**nteroperable, and **R**eusable) principles (Barker et al., 2022; Wilkinson et al., 2016) for computational methods and reproducible analysis across institutions. Nevertheless, significant technical and legal obstacles may prevent centralized data processing using workflows, specifically when datasets originate from heterogeneous sources that cannot be easily or legally combined (Rujano et al., 2024). Researchers may not be permitted to access sensitive data directly or transfer them to their own infrastructure. Furthermore, high-throughput processing requires understanding multiple workflow ecosystems, which can be cumbersome and impractical for scientists focused on data-driven research.

These constraints are most acute in healthcare. Genomic data generated in clinical care increasingly feed research and secondary use, but policy now directs that such data be analysed in place. In England, trusted research environments (TRE) (Alliance & Nhsx, 2021; NHS-England, 2022) are the default route for research access to data. In the European Union, the European Health Data Space Regulation (Parliament, 2025) mandates that secondary use take place only in secure processing environments (SPEs), and that health data access bodies review data download access requests, so that only non-personal, aggregated and fully anonymised data ever leave the environment. Genomic data explicitly fall within this jurisdiction. Together, these constraints call for an execution interface through which validated, versioned workflows can be submitted to a protected site, monitored, audited, and through which only reviewed results are returned via an airlock mechanism.

Rather than operating directly in the compute environment, researchers may define reproducible analysis as workflows that can be moved to the data (Thaldar, 2025; Yuen et al., 2021). A common workflow-engine-agnostic interface can support workflows implemented in different languages and executed by different workflow engines to be submitted and managed across different compute infrastructures. At the same time, research data is expected to follow the FAIR principles (Wilkinson et al., 2016). In an extended federated use case, where sensitive data often cannot be moved, and analyses are instead brought to the data (Thaldar, 2025), these requirements extend beyond the data themselves to the way analyses are executed and documented: a standardised interface across heterogeneous workflow ecosystems relates to interoperability, while a machine-readable record of each workflow run, its parameters, and the execution environment relates to the provenance and reproducibility that underpin reusability. A unified execution interface is therefore not only a matter of portability but also a building block for FAIR-aligned analysis across distributed research infrastructures.

The Global Alliance for Genomics and Health (GA4GH) (Rehm et al., 2021) defines interoperable standards and policy frameworks that enable secure sharing of clinical and genomic data across international borders. To this end, the GA4GH’s Federated Analysis Work Stream (FAWS) currently defines the following specifications: the Task Execution Service (TES) API (Kanitz et al., 2024) for executing individual batch tasks across a large variety of compute infrastructures, the Data Repository Service (DRS) API for standardised data object access, the Tool Registry Service (TRS) API (Yuen et al., 2021) for sharing container-based tools and workflows, and the Workflow Execution Service (WES) API for managing workflow runs in a uniform and abstract way across various compute infrastructures and cloud platforms. The abstract API schema does not restrict the compute infrastructure; it only expects a set of capabilities. Thus, clients can submit workflows to a WES implementation without prior understanding of the underlying workflow engine or the underlying execution backend, which may include High-Performance Computing (HPC), cloud, or other compute environments, as every WES-compliant execution site exposes the same endpoints and data model for workflow management. The actual execution infrastructure remains an implementation-specific concern of the WES service.

Workflow execution via HTTPS is not new: AWS HealthOmics, the Cromwell REST API, the Seqera Platform API, DNAStack, Google Batch API, DNAnexus and Arvados allow a client to submit, monitor, cancel and retrieve workflow runs via its own proprietary API. Reaching these platforms requires one adapter each. In federated settings this becomes costly since the data manager decides which adaptations run at the data site. A neutral interface shifts the integration point to the deployment site. WES is not the only GA4GH execution interface, together with TES they address different purposes. One WES request corresponds to one run, while TES operates at the level of an individual task. WES is thus an interface between the researcher and a site while TES is between an engine and its compute backend. The WES API abstraction gives a client a single vocabulary set across sites, decoupled from the engine chosen internally. Several established platforms, including Galaxy, Toil and the Seqera Platform have added WES endpoints alongside their own native interfaces, which shows that providers see value in a neutral interface even where they maintain their own.

## Methods

### WES is a GA4GH-approved standard

WES was developed by the GA4GH Cloud Work Stream, now the Federated Analysis Work Stream (FAWS). Early pre-release versions (0.1.0 to 0.3.0) were shaped by feedback from GA4GH Driver Projects, including the Human Cell Atlas (Regev et al., 2017). Version 1.0.0 was the first version approved as a GA4GH standard. It was announced at the 6th GA4GH Plenary in Basel in October 2018 (GA4GH, 2018) following a testbed in which workflows from Human Cell Atlas, Australian Genomics, TOPMed and the PCAWG project were run against WES implementations on Microsoft Azure, Amazon Web Services, Google Cloud Platform and Illumina infrastructure.

The specification has since been maintained under the GA4GH Product Development and Approval Process, which uses semantic versioning. Version 1.0.1 (January 2023) was a maintenance release that added a JSON schema for the data model, moved the specification to OpenAPI 3 and aligned the /service-info endpoint with the GA4GH Service Info standard. Version 1.1.0 (September 2023) added the PREEMPTED run state, optional fields for the workflow engine and engine version in a run request, and endpoints to list and retrieve individual tasks within a run. Changes in 1.1.0 are additive, and the features it deprecates will be removed only in a future major release (2.0.0). Clients written for 1.0.0 therefore remain able to submit, monitor and cancel runs on 1.1.0 services.

### WES within the GA4GH ecosystem

WES is one of several GA4GH standards designed to work together. Because WES services implement the GA4GH Service Info standard, they describe themselves in a common format and can be listed and found through a GA4GH Service Registry. Workflows can be referenced from registries that implement TRS, input data can be referenced by DRS identifiers, and individual tasks can be dispatched to compute backends through TES (see "Integration of GA4GH Federated Analysis Work Stream APIs with WES"). WES is used by GA4GH Driver Projects and partner infrastructures, including ICGC ARGO, CanDIG and ELIXIR. In Europe, the 1+Million Genomes initiative lists WES, together with TES and TRS, as recommended standards for data processing in its framework for federated genomic data access (Saunders et al., 2019).

WES is developed within the GA4GH FAWS and each release follows the GA4GH Product Development and Approval Process with semantic versioning (“Development and governance of WES” section in Supplementary).

### Interoperability example between WESkit and Sapporo

In the interoperability example, a researcher submits the same Snakemake workflow to two WES v1.1.0 endpoints: Sapporo (Japan) running in AWS EC2 and WESkit (Germany) running in the de.NBI cloud. The experiment was conducted on two distinct subsets of the 1000 Genomes Project (Genomes Project et al., 2015; Lowy-Gallego et al., 2019) from publicly available data and distributed to both sites. The example workflow compared allele frequencies at three population-differentiated SNPs between two sub-cohorts: Japanese in Tokyo (JPT) (n=104) and Utah residents with Northern/Western European ancestry (CEU) (n=99). Each cohort’s VCF was held locally at dedicated sites.

A researcher submitted the same workflow to the WES endpoints. The example workflow consists of three steps: 1) subsetting local VCFs to specific samples and SNP regions (*bcftools view*) (Danecek et al., 2021), 2) computing allele counts and frequencies (*bcftools + fill-tags* plugin), 3) extracting summary TSV files. Bcftools (version 1.21) was installed in the conda (version 24.11.3) environment alongside Snakemake (version 9.16.3). Output files contained only aggregated summary statistics with no per-sample genotypes. A workflow analysis request was issued as HTTP POST to each endpoint with site-specific parameter files. Run status was polled via the */runs/<RUN_ID>/status* endpoint until each run terminated. Summary TSV files were downloaded from each site and combined using *aggregate.py* that computed odds ratios and Fisher’s exact test p-values for each SNP across both cohorts.

### Collection of WES API implementations

Table 1 includes 13 manually collected WES implementations that have merged at least one commit within the last 12 months. The adopted WES version was identified by inspecting the respective repositories or official documentation in October 2026. Earlier implementations without recent activity, such as those used in the 2018 GA4GH testbed (including those deployed across four cloud and commercial providers), are not captured in Table 1 as it is a snapshot of actively maintained implementations.

**Table 1:** Platforms and standalone tools implementing the WES API and updated within the last 12 months (October 2026).

| Tool | Publication / WES support since | Repository link | WES version | Supported workflow engines | Language | Licence |
| --- | --- | --- | --- | --- | --- | --- |
| WESkit | (Schneider-Lunitz et al., 2026) | https://gitlab.com/one-touch-pipeline/weskit | 1.1.0 | Nextflow, Snakemake | Python | MIT |
| Sapporo | (Suetake et al., 2022) | https://github.com/sapporo-wes/sapporo-service | 1.1.0 | cwltool, Nextflow, Snakemake, ep3, StreamFlow | Python | Apache-2.0 |
| Workflow-service | Not published | https://github.com/common-workflow-language/workflow-service | 1.0.0 | Arvados, Toil | Python | Apache-2.0 |
| conquest | Not published | https://github.com/anuragxxd/conquest | 1.1.0 | Nextflow, WDL, Snakemake | Go | No Licence file |
| metis | Not published | https://github.com/JaeAeich/metis | 1.1.0 | Nextflow | Rust | Apache-2.0 |
| Galaxy platform | release 26.0 | https://galaxyproject.org/ga4gh/#workflow-execution-service-wes-api | 1.0.0 | Galaxy workflows (.ga, gxformat2) | Python | MIT |
| Seqera | https://seqera.io/whitepapers/enabling-collaborative-data-analysis-at-scale-with-seqera-platform/ | https://docs.seqera.io/platform-api/info/ga4gh-info | 1.0.0 | Nextflow | Closed source | Proprietary |
| Toil | release 5.6.0 | https://toil.readthedocs.io/en/master/running/server/wes.html | 1.0.0 | CWL, WDL, Toil native workflows. | Python | Apache-2.0 |
| cromwell-wes-service | Not published | https://github.com/DNAstack/cromwell-wes-service | 1.0.0 | WDL | Java | LGPL-2.1 |
| bento_wes | Not published | https://github.com/bento-platform/bento_wes | 1.1.0 | WDL | Python | LGPL-3.0 |
| GA4GH-WES-API-Service | Not published | https://github.com/NGS360/GA4GH-WES-API-Service | 1.1.0 | CWL, WDL | Python | MIT |
| cwl-WES | Not published | https://github.com/elixir-cloud-aai/cwl-WES | 0.3.0 | CWL | Python | Apache-2.0 |
| proWES | Not published | https://github.com/elixir-cloud-aai/proWES | 1.0.0 | n/a (gateway) | Python | Apache-2.0 |

## Results

WES is one of the APIs developed by the GA4GH FAWS. Together, they define interfaces for discovering, accessing, and processing biomedical data across different compute environments. A diverse community of bioinformaticians, computational scientists and infrastructure providers is developing WES, each with different expectations for workflow execution. The following principles guide the WES design.

1. Researchers should primarily interact with a simple high-level interface rather than engine-specific controls.
2. Workflow execution should capture sufficient information about each run (workflow, version, parameters, engine, etc.) to support reproducing it in a different compute site.
3. WES should integrate with established security mechanisms, such as OAuth 2.0 bearer tokens and GA4GH Passports, without prescribing a specific authentication/authorisation model.
4. WES should support workflow execution across a wide range of deployment environments, from small- to large-scale infrastructures supporting analysis of various cohorts.

To address these needs, WES defines a minimal interface to submit, monitor, and manage workflow runs uniformly. By abstracting away workflow-engine and compute-infrastructure specifics, as well as execution scheduling, the WES specification lets users focus on the analysis itself rather than the operational details. Furthermore, when deployed at the site where the data reside and combined with the GA4GH DRS API for data resolution, the WES API can simplify data logistics by bringing the computation to the data. Logs defined by the WES data model allow researchers to inspect workflow execution outcomes, including failure states, thereby facilitating targeted debugging and optimization.

### Data Model

The GA4GH WES API (v1.1.0) organises workflow execution around a set of components that capture the workflow lifecycle. A client initiates a workflow execution by submitting a *RunRequest* that specifies the workflow, input parameters and optional settings.

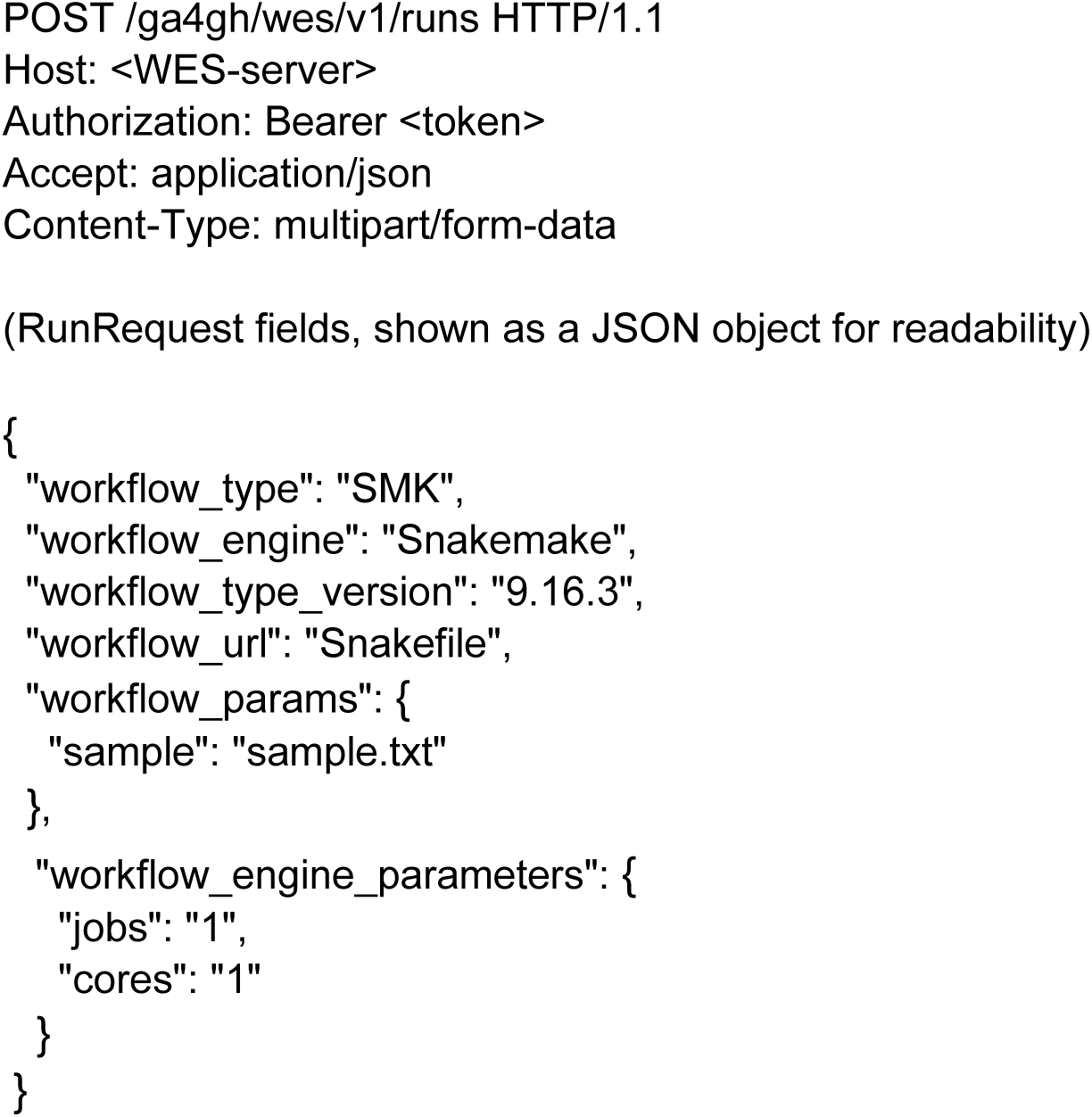

Please note that *workflow_type_version* is the *language* version. For Snakemake and Nextflow the *workflow_type_version* happens to equal the engine version.

The WES implementation creates a new *RUN* and returns its corresponding *RunID.* A *Run* transitions through several execution states, non-terminal: UNKNOWN, QUEUED, INITIALIZING, RUNNING, PAUSED, CANCELING, terminal: COMPLETE, EXECUTOR_ERROR, SYSTEM_ERROR, CANCELED, PREEMPTED (added in v1.1.0). Clients monitor progress through the *RunStatus* object. Although the 11 states can be grouped into non-terminal and terminal states, the specification does not define a transition model. It is up to the implementer to map engine states onto this model. While the Cromwell engine provides job states, Nextflow and Snakemake report progress through logs and exit codes. Each *Run* is associated with a *RunLog* component that records the original execution request, execution-state, workflow logs and output after completion. General information about the WES server can be obtained via the *ServiceInfo* component.

### Interface

The REST endpoints (Figure 1, Supp. Table S1) exposed by the WES API uses the WES data model components to control the workflow run lifecycle. A client first submits a *RunRequest* using POST /runs, which creates a new Run and returns a *run_id.* This id is used to query the workflow status (GET /runs/<*run_id*>/status) or cancel an active workflow (POST /runs/<*run_id*>/cancel). Additional endpoints allow retrieval of service information (GET /service-info) and listing of all workflow runs (GET /runs). The API lets clients interact with a WES implementation through a uniform interface independent of the applied workflow engine or compute infrastructure. Example requests and the corresponding responses are shown in Supp. Table S2.

**Figure 1:**
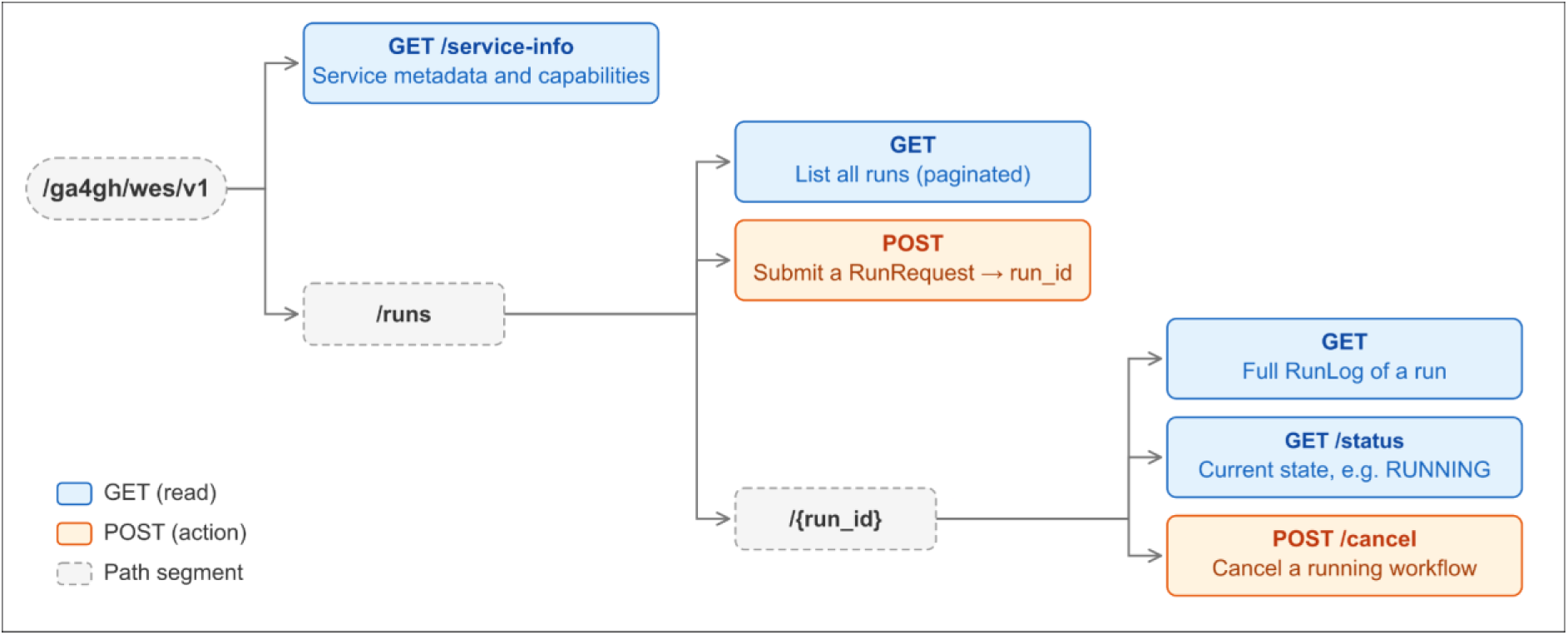
Endpoints of the GA4GH WES API (v1.1.0). The two task-level endpoints (GET /runs/{run_id}/tasks and GET /runs/{run_id}/tasks/{task_id}) are not shown. Expected HTTP status codes are 200 (success) and 400, 401, 403, 404 and 500 (errors).

A full description of the WES API can be found at: https://ga4gh.github.io/workflow-execution-service-schemas/docs/

### Use cases and Application

WES supports single-user deployment, in which a client submits execution requests directly to a locally deployed WES server, as used for development, testing or on-premises clinical settings (Supp. Figure S1A), and multi-user deployments, in which one WES instance dispatches runs to several compute infrastructure, for example in federated or cloud environments (Supp. Figure S1B).

### Implementations

The WES standard has been adopted by a range of open-source projects and a commercial platform. We have collected 13 adoptions of the WES API, which can be grouped into three categories (Table 1). Six are standalone services that expose the WES API over multiple workflow engines: WESkit (Snakemake, Nextflow), Sapporo (cwltool, Nextflow, Snakemake, ep3, StreamFlow), workflow-service (Arvados, Toil), conquest (Go; Nextflow, WDL and Snakemake on Kubernetes), metis (Nextflow) and cwl-WES (CWL). Six further adoptions are workflow systems or analysis platforms that expose the WES API in addition to their native interfaces: Galaxy, which serves WES 1.0.0 as of release 26.0, the Seqera Platform, which accepts WES run requests for Nextflow workflows, Toil, which provides an experimental WES API implementation, cromwell-wes-service, a wrapper for the Cromwell workflow engine developed by DNAStack, bento_wes by the Bento platform and the GA4GH-WES-API-Service. Finally, proWES does not execute run requests but rather acts as a gateway that injects middleware, for example to route requests to configured WES servers.

### Integration of GA4GH Federated Analysis Work Stream APIs with WES

By integrating with the broader suite of GA4GH FAWS APIs, WES enables a “search-access-analyse” process where analytical methods are directly brought to the data’s location, reducing the technical barriers of centralized data pooling and helping clinicians and researchers comply with constraints on data movement. In combination with other FAWS APIs, WES can use TRS (Yuen et al., 2021) for discovery of the standardised workflow definition and containerized tools, and DRS to map logical identifiers to physical data objects independent of the underlying systems (Figure 2). For example, a WES run request can reference a workflow hosted in a TRS repository through a *workflow_url*, while the workflow inputs can be specified using DRS URIs (drs://…) that identify the required data objects. WES can then retrieve or resolve the workflow definition and associated tools through TRS and resolve the referenced data objects through DRS before executing the workflow. This composition is architectural. Among the thirteen adoptions surveyed, platforms like Toil, Galaxy and Cromwell resolve TRS or DRS identifiers server side. WESkit is actively working on TRS and DRS integration, although this remains experimental as of October 2026.

**Figure 2:**
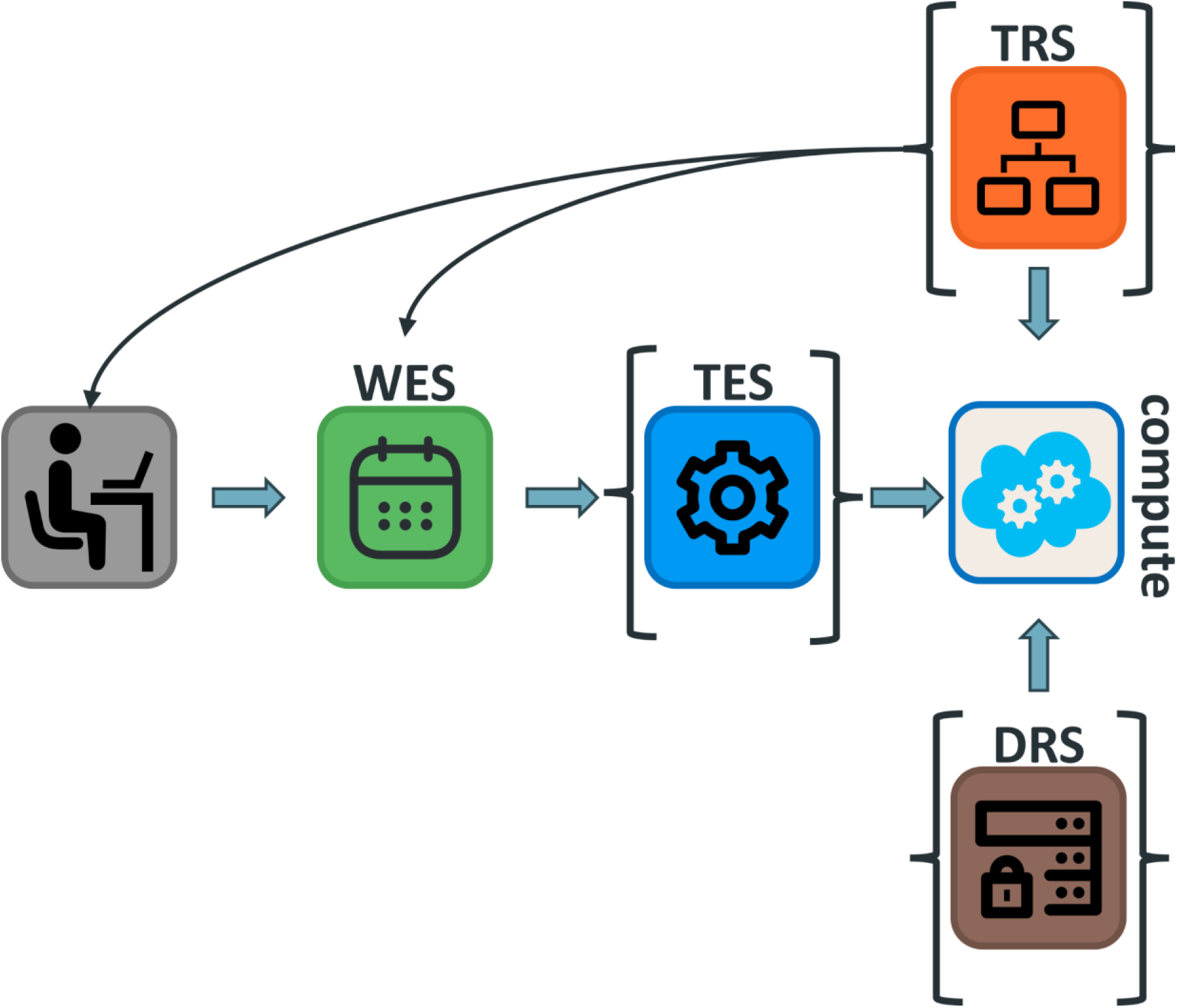
In a comprehensive deployment scenario, the client/user initiates execution through WES, workflows and tools may be discovered and retrieved via TRS at different stages, TES may be used for task-level orchestration and execution, and DRS provides access to data at the compute backend. TES serves as an optional intermediary layer, and neither TRS nor DRS is strictly required in all deployment configurations.

This separation allows a WES service to orchestrate analysis without requiring the caller to know the physical location of the workflow tools or input data. Furthermore, WES can leverage TES (Kanitz et al., 2024), a GA4GH API for submitting, managing, and executing computational tasks, to send individual workflow tasks to diverse compute backends, such as cloud platforms or HPC queuing systems. Ultimately, this abstraction shields researchers from infrastructure-specific complexities, ensuring that computational workflows can be executed portably and documented consistently across distributed research infrastructures, which supports, but does not by itself, ensure their FAIRness.

### Interoperability example between WESkit and Sapporo

The WES API enables workflow request abstraction between implementations as showcased by a demonstration workflow [https://doi.org/10.5281/zenodo.21184328] that runs a federated genomic analysis across two sites (Figure 3). Each site hosts a different population subset of the 1000 Genomes cohort and serves it through a different WES implementation: Sapporo at one site and WESkit at the other.

**Figure 3:**
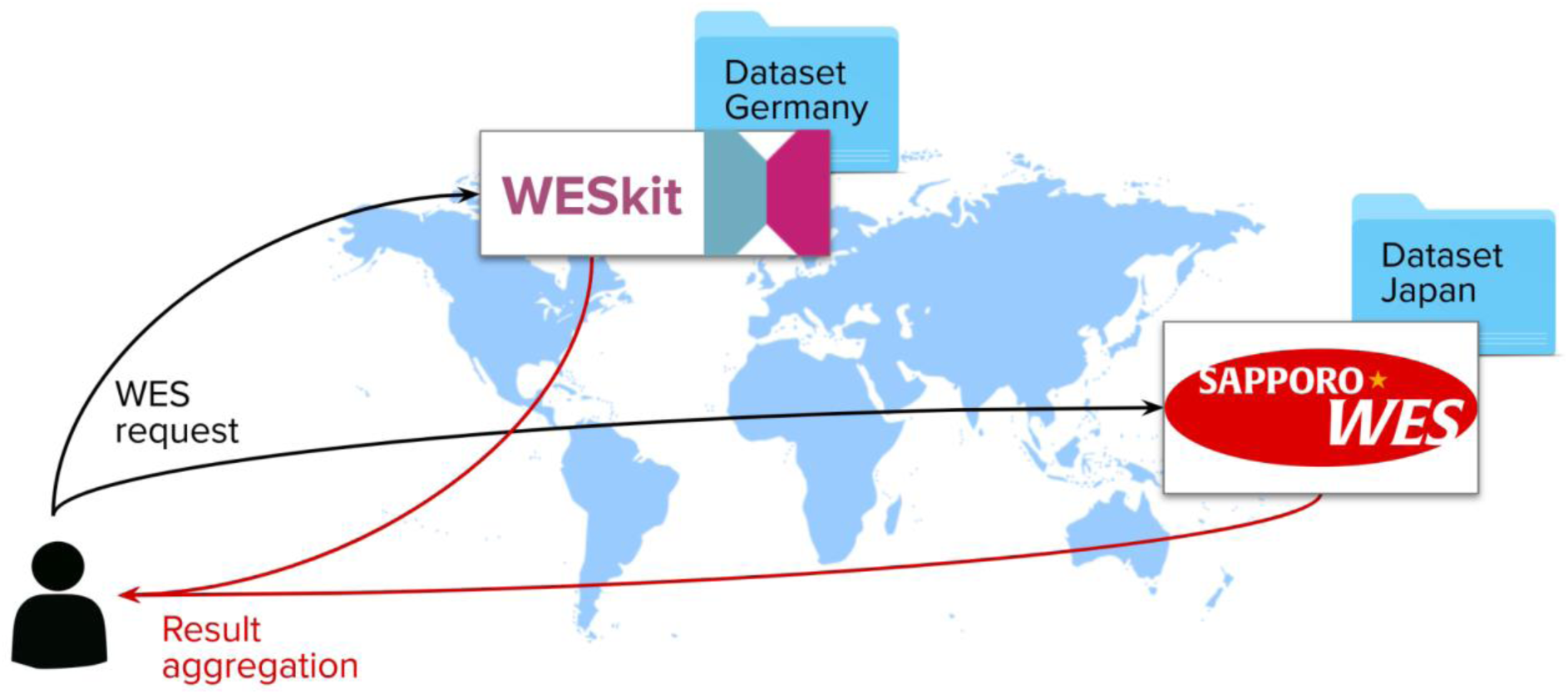
Federated genomic analysis using two WES servers. Each site has its own subset of a large cohort. A researcher without direct access to the data runs a workflow via WES at both sites to receive and aggregate the results.

The WES API allows both sites to understand the execution request through standardised HTTPS request and REST semantics. The individual-level genotype data (VCF) is treated as sensitive and does not leave either site. Only per-site allele frequencies are returned, which are then combined into the final result. However, for different use-cases, the submission would require minor modifications of the request, as functionalities such as downloading workflow files and parameter provisioning are implemented differently by Sapporo and WESkit respectively (Supp. Table S3).

### Security and access control

The specification addresses authentication at the transport level and leaves the authorization to the implementations. Credentials are passed in the HTTP Authorization header, and a service advertises how to obtain them through auth_instructions_url in /service-info. The specification does not define identity provider, scope or authorization decision. GA4GH Passports (Voisin et al., 2021) define how this information is provided but are not required by WES. An executed code via WES may attach workflow files, a remote workflow_url, and arbitrary workflow engine commands which deployments need to take care of. Any restriction is non-standard and is specific to the respective implementation.

### WES and the FAIR principles

WES contributes to FAIR analysis primarily through provenance and execution interoperability rather than data description. Following the FAIR principles for computational workflows (Wilkinson et al., 2025) and the provenance requirement of the original data principles (R1.2), each WES run produces an engine-independent, machine-readable *RunLog* that remains retrievable for as long as the service retains it. It contains information about workflow location, type and version, engine and engine version, parameters, and tags, together with the current state, the executed command, exit code and logs. This record documents how a given result was generated in a form identical across workflow engine executions and compute backends, which is the precondition for reproducing or re-running an analysis without knowledge of the executing site. The specification deliberately delegates workflow and data findability and accessibility to TRS and DRS and does not define persistent identifiers for runs or outputs, output checksums, or a structured description of the execution environment. Our two-site demonstration illustrates both sides: in this case, the run records were sufficient to reproduce the analysis, whereas differences in parameter provisioning between implementations point to the parts of the interface that are not yet fully specified. As next steps toward FAIR-compliant workflow execution, we put forward three extensions for discussion in the WES community: an optional provenance export, a minimal contract for output description, and globally resolvable run identifiers.

## Discussion

We have described the GA4GH WES API and its usage in the community. We identified 13 adoptions of the specification: 12 services that execute workflows and one gateway (proWES) that routes requests and allows middleware to be injected. These adoptions span the open-source community and commercial providers. In our showcase, run requests with an identical structure, but with site-specific adjustments, executed the same analysis at two independently developed WES services in Germany and Japan without transferring individual-level genotype data at analysis time, demonstrating the abstraction capabilities of the WES API. The shared vocabulary for endpoints, run states, and the structure of the request was enough for two WES services to accept and execute a run. Still, input provisioning and output retrieval differed at each site and results had to be downloaded through non-standard endpoints. The specification establishes a common submission and monitoring vocabulary while the undefined parts show where implementations diverge.

The breadth of adoption suggests that the specification is inexpensive to implement; WES has been reimplemented in Python, Java, Go, and Rust, by individual developers, small teams and platforms alike, which is feasible because the specification comprises eight endpoints and a small data model. The split between stand-alone and platform implementations is equally informative. Established platforms such as Galaxy, Toil and the Seqera Platform added WES endpoints alongside their mature native interfaces, indicating that the standard’s value lies in providing an entry point for workflow execution rather than in replacing engine-specific tooling. Implementations target one of two specification versions: workflow-service, Galaxy, Seqera, Toil, cromwell-wes-service, proWES targeting v1.0.0 whereas WESkit, Sapporo, GA4GH-WES-API-Service, conquest, metis, and bento_wes implement 1.1.0. Because v1.1.0 adds endpoints and optional fields without removing any, and its deprecations take effect only in v2.0.0, core operations such as submitting, monitoring, and cancelling runs should work across both versions. Finally, cwl-WES adopts version 0.3.0 and even though we identified a merged commit within the last 12 months it was just an update on the Docker file rather than updating the WES specification. We did not complete a full conformance assessment for all implementations. Table 1 should therefore be read as a record of WES API adoptions, and a conformance suite, building on the 2018 GA4GH WES testbed, would close this gap and is a priority for the Federated Analysis Work Stream. General-purpose clients exist, such as the WES client in the Dockstore command-line tool, which supports bearer-token and AWS authentication, and wes-client from the workflow-service project. However, many implementations provide their own command-line client, and there is no reference client library maintained alongside the specification. The WESkit client (https://pypi.org/project/weskit-client), for example, exposes additional routes that are not part of the specification.

The interoperability example showed that under the same conditions: a researcher without access to either dataset obtained allele frequencies at three SNP positions from both sites using the same workflow and the same request structure, while the variant calls themselves were processed locally and never left their site. This demonstrates a federated analysis executed between WESkit and Sapporo. It is nonetheless a small feasibility example, involving two implementations, one workflow language, three variants and two sub-cohorts totaling 203 individuals. One source of friction remains: the run requests require site-specific adjustments, as both WESkit and Sapporo provide files and parameters differently. The WES specification does not dictate these mechanisms, so implementers can choose the strategy that suits their infrastructure. However, a run request must be portable across implementations to the extent that both accept the same input format. The demonstration was designed to test interoperability of the API, not engine abstraction or governance, and has several limitations. Both sites ran Snakemake, so no engine abstraction was shown. The executed workflow was pre-installed at WESkit but shipped with the request to Sapporo, so workflow portability was shown at one endpoint only and the results were retrieved through a custom download endpoint (Supp. Table S3). The public data input was pre-staged to emulate a federated analysis rather than encountering governance and neither authentication, Passports, TRS, DRS nor TES were applied. Running a second language at one site would have addressed the engine abstraction limitation.

WES gains much of its value by combining other GA4GH FAWS APIs, TRS for workflow and tool discovery, DRS for resolving data identifiers to physical objects, and TES for dispatching individual tasks to compute backends. Resolving DRS and TRS references at execution time requires handling signed URLs and GA4GH Passports (Voisin et al., 2021), which the WES specification does not yet address, and is important for general federated analysis beyond the read-only case demonstrated here. This issue has a concrete consequence for healthcare deployments. TREs do not release results automatically; rather, they require output checking by trained staff for each egress request. Output checking typically involves two trained reviewers per request, which will likely be a bottleneck. Because WES 1.1.0 does not specify outputs, we propose that the workflow declares what outputs will be produced and the service exposes only those objects. A review then needs to focus on the comparison of what was declared and what was actually produced rather than inspection of arbitrary files. This would give secure processing environments a standard basis for output checking. In the United States, NIH policy (NOT-OD-24-157, effective 25 January 2025) requires that systems holding controlled-access genomic data meet NIST SP 800-171 (Ross & Pillitteri, 2024). In the European Union, Article 73 of the European Health Data Space (Parliament, 2025) regulation obliges secure processing environments (SPEs) to restrict access, retain identifiable logs and undergo regular audits. Detailed requirements for these environments are due in Commission implementing acts by March 2027. This is not something the WES specification should take care of but rather support them by carrying credentials in the standard header, letting the service provide authentication and leave the authorization to the site.

Three specification extensions follow directly from the FAIR gaps identified above. We present them as proposals for the WES community to take forward through the GA4GH PDAP minor-release track, each tracked as an issue on the specification repository. First, an optional provenance export (GET /runs/<RUN_ID>/provenance) aligned with Workflow Run RO-Crate (Leo et al., 2024) would close the corresponding reusability gap. The *RunLog* already includes information such as parameters, engine version, and run logs in a machine-readable format, but this information is retrievable only from the issuing service and only while it retains it. Delivering this object matters because the RO-Crate must outlive the service. Sapporo already generates RO-Crate metadata which confirms feasibility. Second, the WES specification does not define the output format. Thus, a minimal contract for output description (GET /runs/<RUN_ID>/outputs) would define what a service must report: at least a stable identifier, a name or path, media type, a size, and a checksum. Having this, a client could verify a run’s output without implementation-specific knowledge. Third, a globally resolvable run_id would address the FAIR criterion (F1) that is currently delegated to external services. A run_id is unique within one instance, so this run_id cannot be resolved or deduplicated across sites, a situation that federated use cases create by design. Beyond the three FAIR-driven extensions, the community requires additional run lifecycle endpoints for cleaning and deleting runs, and richer structured logging. Given the heterogeneity of workflow engines and their non-standardised log formats, making harmonised status reporting, rather than log-parsing, is a near-term goal. Each change should be introduced under clear versioning and should preserve backwards compatibility.

The GA4GH WES API shows that a minimal engine-agnostic specification can support moving workflows and data across institutions and jurisdictions, within each site’s governance constraints. The remaining work should focus on specifying its edges precisely enough that independent implementations agree, strengthening federated analysis before extending the interface.

### Future directions

A further argument for a constrained execution interface is emerging from autonomous agents. Language-model agents are increasingly given direct access to domain APIs, GeneGPT (Jin et al., 2024) for example answers genomic questions by invoking NCBI Web APIs. Granting agents general-purpose access to compute is difficult to audit, secure and reproduce but raise work on making agent actions observable (Chan et al., 2024). An agent restricted to submitting WES requests that bind TRS-registered workflows to DRS-identified inputs would produce submissions that are explicit and auditable, because each request would be expressed through a small set of OpenAPI-defined operations. Depending on the execution-site policy, the APIs provide a well-defined boundary where access control, audit, and human oversight can be applied, and allow an autonomous AI tool to operate across federated sites on a common interface. The GA4GH has begun to explore this through an experimental Agentic Harness Profile, which is not a GA4GH-approved standard, and the GA4GH AI Work Stream has identified governance, provenance and benchmarking as priority areas for standards in this space.

## Conclusion

The described work presents the GA4GH WES API, a standard that allows submission of workflow execution via HTTP requests. Its abstract design allows users to submit standardised execution requests without any prior knowledge of the underlying compute infrastructure. WES has been adopted by 13 open-source and commercial services and recommended by the 1+Million Genomes framework, which shows a clear need for standardised, HTTP-based workflow execution across compute infrastructures. A conformance suite and the extensions proposed here are the next priorities for the GA4GH Federated Analysis Work Stream.

### Data Availability

The reported GA4GH WES API describes a standard and did not generate any new primary data.

The demonstration analysis reported in section ”Interoperability example between WESkit and Sapporo” used a publicly available dataset from the 1000 Genomes Project (Genomes Project et al., 2015) from the International Genome Sample Resource (https://ftp.1000genomes.ebi.ac.uk/vol1/ftp/data_collections/1000_genomes_project/).

The demonstration workflow, its configuration, the WES request submitted to each endpoint, and the aggregation script are archived at https://doi.org/10.5281/zenodo.21184328.

### Code Availability

The GA4GH Workflow Execution Service (WES) (v1.1.0, Apache Licence 2.0) definition can be found at: https://github.com/ga4gh/workflow-execution-service-schemas

WESkit (v1.0.0) can be found at: https://gitlab.com/one-touch-pipeline/weskit

Sapporo (v2.2.6) can be found at: https://github.com/sapporo-wes/sapporo-service

### Author Contributions (CRediT)

Valentin Schneider-Lunitz (Conceptualization [lead], Investigation [lead], Visualization [supporting], Writing – original draft [lead]), Landfried Kraatz (Conceptualization [lead], Investigation [lead], Visualization [lead], Writing – original draft [supporting]), Patrick Magee (Conceptualization [supporting], Investigation [lead], Writing – original draft [supporting]), Venkat S. Malladi (Investigation [lead], Visualization [supporting], Writing – original draft [supporting]), Brian D. O’Connor (Investigation [lead], Visualization [supporting], Writing – original draft [supporting]), Susheel Varma (Conceptualization [lead], Investigation [lead], Supervision [lead], Funding acquisition [lead], Writing – review & editing [supporting]), James Eddy (Investigation [lead]), Hirotaka Suetake (Investigation [supporting]), Jeff Gentry (Investigation [supporting]), Michael R. Crusoe (Investigation [supporting]), Angel Pizarro (Investigation [supporting]), Junjun Zhang (Investigation [supporting]), Denis Yuen (Investigation [supporting]), Kyle Ellrott (Investigation [lead]), Philip R. Kensche (Investigation [supporting]), Michael Lukowski (Investigation [supporting]), Reggan Thomas (Conceptualization[supporting]), Sasha V. Siegel (Conceptualization[supporting], Writing - review & editing [supporting] ), Alexander Kanitz (Investigation [lead], Visualization [supporting], Writing – original draft [supporting]), Tazro Ohta (Conceptualization [lead], Investigation [lead], Writing – original draft [supporting]), Sven O. Twardziok (Supervision [lead], Funding acquisition [lead], Conceptualization [lead], Investigation [supporting], Writing – original draft [supporting])

### Conflict of interests

None declared.

## Funding

The Global Alliance for Genomics and Health (GA4GH) receives funding from the National Human Genome Research Institute, with co-funding from the National Cancer Institute (NCI), the National Heart, Lung, and Blood Institute, and the Office of Data Science Strategy under award number U24HG011025; the Canadian Institutes of Health Research (CIHR) (204310); the Wellcome Trust (220544/Z/20/Z); the UK Medical Research Council (MC_PC_19024); the Ontario Institute for Cancer Research; the Broad Institute of MIT and Harvard; and the European Molecular Biology Laboratory (EMBL).

This work received funding by ELIXIR Europe, the research infrastructure for life science data through ELIXIR Travel Grant and ELIXIR Biohackathon Europe and the ELIXIR Compute Platform.

The project was funded by the Federal Ministry of Research, Technology and Space (Bundesministerium für Forschung, Technologie und Raumfahrt, BMFTR) as part of the German Center for Child and Adolescent Health (under the funding code 01GL2401A).

This work was funded by the Federal Ministry of Research, Technology and Space (Bundesministerium für Forschung, Technologie und Raumfahrt, BMFTR) in the frame of de.NBI/ELIXIR-DE (W-de.NBI-001).

This work has been funded by ROIS-DS-JOINT (039RP2024) and the SECOM Science and Technology Foundation through the SECOM Challenge 2023 Research Grant

This work was funded by the NIH NCATS grant #U24TR002306

## Acknowledgement

We thank the members of the GA4GH Federated Analysis Work Stream that built the WES GA4GH standard, and the GA4GH Driver Projects that tested WES, including Human Cell Atlas, Australian Genomics and TOPMed. Peter Isaac Amstutz, David Steinberg, David Glazer, Abraham Chavez, Alex Strizhak, and Ania Niewielska were involved in the start of the Federated Analysis Work Stream (formerly Cloud WS) and made significant contributions to the WES API.

This work was supported by the de.NBI Cloud within the German Network for Bioinformatics Infrastructure (de.NBI) and ELIXIR-DE (Forschungszentrum Jülich and W-de.NBI-001, W-de.NBI-004, W-de.NBI-008, W-de.NBI-010, W-de.NBI-013, W-de.NBI-014, W-de.NBI-016, W-de.NBI-022).

Claude (Anthropic) was used to assist with language and consistency checking. Suggestions were reviewed and approved by authors.

## Notes

### Competing Interest Statement

The authors have declared no competing interest.

