## Supplementary Data for "The GA4GH Workflow Execution Service: A Standard API for Workflow Execution and Management in Genomics"

Authors (as in main text)

References are listed in the main text

### Supplementary Note S1: Development and governance of WES

The development of the Federated Analysis Work Stream APIs is governed by regular meetings of all API co-leads where current progress is presented and open questions are discussed. Proposed features are presented and reviewed by work stream members with relevant expertise and by interested members of the wider community. The community discusses requests for their potential use, considering the potential use for the whole community, implementation effort and effect on the API’s structure. Implementers are welcome to present their results and ideas.

The WES standard is refined gradually by the community. New features and modifications are requested by implementers and users via issues on GitHub. Contributors may create pull requests (PR) to propose code changes. The review process differs significantly from typical software development projects. It is a GA4GH specification and thus the review is a consensus-driven process. Invited reviewers and project leads review and comment on the PR. Changes are discussed until the Work Stream reaches consensus. Because WES is an approved GA4GH product, new releases follow the GA4GH Product Development and Approval Process (PDAP) (GA4GH, <https://ga4gh.org/our-products/development-and-approval-process>). Before approval, a product must have at least two independent implementations, at least one of them open source. Approval follows a public comment period of at least one month, review by the Regulatory & Ethics and Data Security Work Streams and by a Product Review Committee, which also checks alignment with the recommendations of the Technical Alignment Subcommittee (TASC), and a vote of the GA4GH Product Steering Committee. After approval, changes follow one of three tracks: patch releases for urgent fixes, minor releases for continued development, and major releases for changes of scope or breaking changes. Upon consensus, the PR is merged into the “develop” branch, and GitHub Pages automatically deploys a preview of the updated documentation. When enough changes are collected, a release branch is created together with a new WES API version. The release branch is merged into “develop” and “master” and a new tag version is created. This consensus-based process is slower than typical software development, but it is what makes a GA4GH standard stable enough for national and clinical infrastructures to rely on, as in other standards development organisations.

### Supplementary Note S2 API reference

| Endpoint URI | Type | Description |
| --- | --- | --- |
| /service-info | GET | General information about the service: authentication endpoints, supported workflow types, supported engines, and maintainer contact information. |
| /runs | GET | Listing all workflow runs (ongoing and finished; paginated). |
| /runs | POST | Submit a new workflow execution. Returns a RUN_ID for that run. |
| /runs/<run_id> | GET | Detailed information about a given workflow execution. |
| /runs/<run_id>/status | GET | Status of the workflow execution, e.g. RUNNING, SYSTEM_ERROR. |
| /runs/<run_id>/tasks | GET | List of tasks executed for a given workflow. |
| /runs/<run_id>/tasks/  <task_id> | GET | Detailed information about a given task. |
| /runs/<run_id>/cancel | POST | Cancel a running workflow. |

**Supp. Table S1:** WES API v1.1.0 endpoints. Expected HTTP status codes are 200 (success) and 400, 401, 403, 404 and 500 (errors).
A full description of the WES API can be found at:
<https://ga4gh.github.io/workflow-execution-service-schemas/docs/>

| Request | Response (200) |
| --- | --- |
| GET /service-info | {"id": "org.ga4gh.myservice", "supported_wes_versions": ["1.1.0"], "workflow_engine_versions": {"Snakemake": {"workflow_engine_version": ["9.16.3"]}} , "system_state_counts": {"RUNNING": 2, "COMPLETE": 17}, …} |
| POST /runs | {"run_id": "b3f1c2a8"} |
| GET /runs | {"runs": [{"run_id": "b3f1c2a8", "state": "COMPLETE"}, {"run_id": "9de04f17", "state": "COMPLETE"}]} |
| GET /runs/b3f1c2a8 | {"run_id": "b3f1c2a8", "state": "COMPLETE", "run_log": {"name": "...", "cmd": [...], "start_time": "...", "end_time": "...", "stdout": "...", "stderr": "...", "exit_code": 0}, "outputs": {…}} |

**Supp. Table S2**: Example WES requests and abbreviated JSON responses.

### Supplementary Note S3 Deployment scenarios

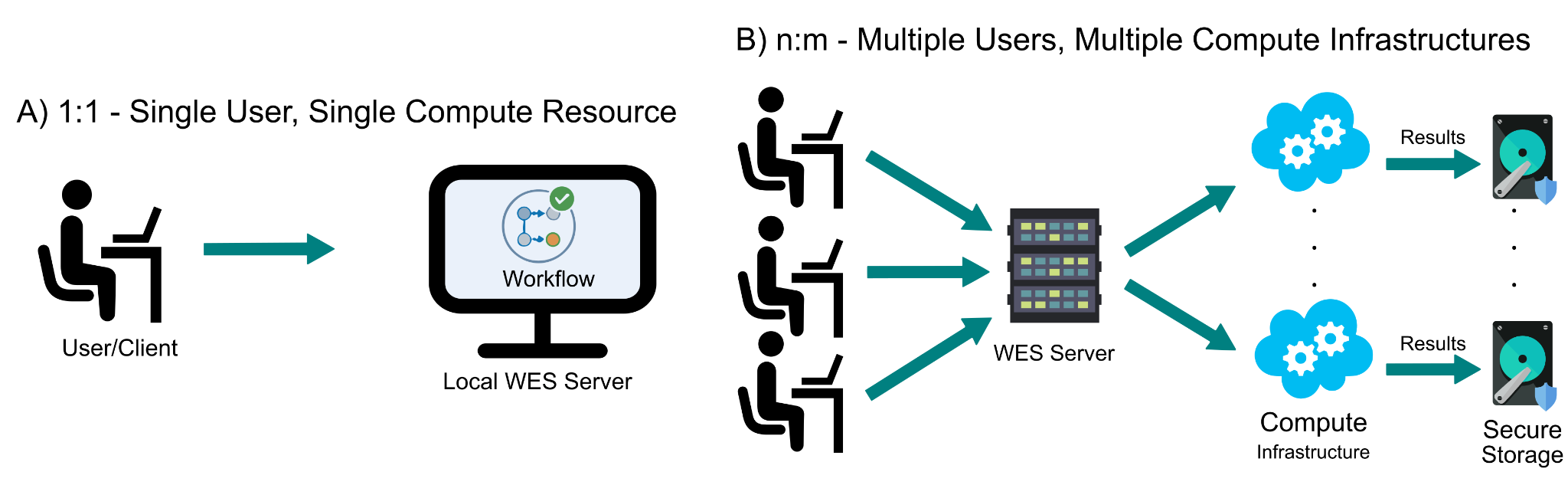
**Supp. Figure S1: 1:1 and n:m deployments.** (A) 1:1 deployment scenario in which a single client submits a workflow execution request to a locally deployed WES instance operating on a dedicated compute resource. (B) An *n:m* deployment scenario in which multiple clients submit workflow requests to a WES server, which routes workflows to predefined compute infrastructures. Results are written to storage systems managed by the respective compute infrastructure.

### Supplementary Note S4 Federated demonstration

|  | Sapporo | WESkit |
| --- | --- | --- |
| workflow_url | Remote URL or name of an attached file. | Relative ‘*file:*’ path, resolved against attachments of pre-installed workflows |
| Input data | Host directory mounted into the engine container. | Worker volume mounted paths. |
| Output retrieval | Non-standard GET endpoint. | Non-standard GET endpoint. |
| Output provenance (via *RunLog* object) | Mapping of *file:url* objects. | A flat list of every file in the run directory. |

**Supp. Table S3**: Implementation differences between Sapporo and WESkit relevant to the federated run.

These differences are possible because the WES API does not explicitly specify their implementation, with the intention of letting implementers choose the best-fitting approach for their use case.
